# Neural network modelling reveals changes in directional connectivity between cortical and hypothalamic regions in obesity

**DOI:** 10.1101/2020.05.10.087619

**Authors:** Katharina Voigt, Adeel Razi, Ian H. Harding, Zane B. Andrews, Antonio Verdejo-Garcia

**Affiliations:** School of Psychological Sciences and Turner Institute for Brain and Mental Health, Monash University, Victoria, Australia; The Wellcome Centre for Human Neuroimaging, University College London, London, WC1E 6BT, UK; Department of Electronic Engineering, NED University of Engineering and Technology, Karachi, Sindh, 75270, Pakistan; Biomedicine Discovery Institute and Department of Physiology, Monash University, Victoria, Australia

## Abstract

Obesity has been ascribed to corticostriatal regions taking control over homeostatic areas. To test this assumption, we applied an effective connectivity approach to reveal the direction of information flow between brain regions and the valence of connections (excitatory versus inhibitory) as a function of adiposity and homeostatic state. Forty-one participants (21 overweight/obese) underwent two resting-state fMRI scans: after overnight fasting (hunger) and following a standardised meal (satiety). We used spectral dynamic causal modelling to unravel hunger and adiposity related changes in directed connectivity between cortical, insular, striatal and hypothalamic regions. During hunger, as compared to satiety, we found increased excitation of the ventromedial prefrontal cortex over the ventral striatum and hypothalamus, suggesting enhanced top-down modulation compensating energy depletion. Adiposity was associated with increased excitation of the anterior insula over the hypothalamus across the hunger and satiety conditions. The interaction of hunger and adiposity yielded decreased intra-cortical excitation from the dorsolateral to the ventromedial prefrontal cortex. Our findings suggest that obesity is associated with persistent top-down excitation of the hypothalamus, regardless of homeostatic state, and hunger-related reductions of dorsolateral to ventromedial prefrontal inputs. These findings are compatible with eating without hunger and reduced self-regulation views of obesity.

**Significance Statement:** Obesity is a leading contributor to morbidity and mortality. Neurobiological theories propose that, in obese people, corticostriatal regions take over homeostatic areas. Neuroimaging-based functional connectivity is well-poised to unravel such abnormalities by examining between-regions communication, but existing studies have only measured signal covariance, not direction and valence of connectivity. We applied computational modelling to reveal the direction of information flow between brain regions and excitatory/inhibitory valence of connections in obese versus healthy-weight participants. Obesity associated with heightened top-down excitation from the insula to hypothalamus, and reduced excitation within prefrontal regions. Findings have two advantages relative to current knowledge: demonstrate theory-based directional abnormalities, i.e. cortical regions taking over homeostatic areas; and inform brain stimulation therapies targeting cortical input to lower-level regions.

## Introduction

Obesity contributes to ~5% of deaths globally, reduces life expectancy by almost a decade (Grover et al., 2015), and accounts for a 2.8% loss to the global gross domestic product (Tremmel et al., 2017). The prevalence of obesity has risen in parallel with increased access to energy-dense, highly attractive foods (Crino et al., 2015). These food choices can overtake the neural mechanisms that regulate homeostatic-based eating and promote overconsumption (Berthoud, 2012). Neurobiological theories suggest that obesity is underpinned by abnormal interactions between homeostatic and non-homeostatic brain systems: (i) the “hedonic eating” model proposes that the neural regions that code reward valuation (e.g. striatum, medial prefrontal cortex) take over those involved in energy homeostasis (e.g. hypothalamus) (Berthoud et al., 2017; Cameron et al., 2017); (ii) the “eating without hunger” framework similarly suggests that the neural regions that process salient external stimuli outweigh those involved in internal energy sensing (Carnell et al., 2013); and (iii) the “self-regulation” view emphasizes impaired top-down regulation of lower-level striatal and limbic regions (Carter et al., 2016; Volkow et al., 2017). Thus, understanding the interaction between the neural systems pinpointed by these theories is key to envisage new approaches to prevent and treat obesity.

Neuroimaging functional connectivity studies have the potential to improve our understanding of the interplay between homeostatic and non-homeostatic neural systems. Homeostatic states (hunger versus satiety) and body mass index (BMI) have been associated with different strengths of functional connectivity between the hypothalamus, striatum, insula and prefrontal cortex regions (Morton et al., 2014; Liu and Borgland, 2015; Cassidy and Tong, 2017; Livneh et al., 2017; Rossi and Stuber, 2018). Specifically, hunger, as compared to satiety, has been associated with increased resting-state connectivity between the insula, the posterior cingulate cortex and the ventromedial prefrontal cortex (Wright et al., 2016; Al-Zubaidi et al., 2019). Furthermore, BMI has been associated with increased connectivity between the hypothalamus and the striatum, the insula and the ventromedial prefrontal cortex at rest (Kilpatrick et al., 2014; Kullmann et al., 2014; Lips et al., 2014; Wijngaarden et al., 2015), and reduced connectivity between the prefrontal cortex and both the insula and the striatum during food valuation and choice tasks (Verdejo-Román et al., 2017; Harding et al., 2018). However, a critical limitation of all these studies is that they cannot speak to the direction of information flow between the brain regions (e.g., whether they reflect bottom-up or top-down communication) or the valence of the connections (whether they are excitatory or inhibitory). Such information is needed to reveal if/how, as suggested by neurobiological theories, non-homeostatic brain systems take over homeostatic areas.

Here, we applied a novel hypothesis-testing framework to unravel the direction and valence of connections between hypothalamic, striatal, insula and cortical regions as a function of BMI and energy homeostasis. We capitalised on recent advances in modelling the interactions of low-frequency endogenous fluctuations in rsfMRI data using spectral dynamic causal modelling (spDCM; Friston et al. 2014; Razi et al. 2015; Park et al. 2018). In contrast to conventional functional connectivity, spDCM predicts directional communications among distributed brain regions (i.e., effective connectivity; Friston, Harrison, & Penny, 2003). We selected a set of regions featured in neurobiological theories of obesity that showed significant task-evoked activation in response to energy-dense food choices in the same sample that we use here (Harding et al., 2018) (Figure 1A, Table 1). We first used the selected regions to define and estimate a fully connected DCM for each participant (i.e., all ROIs connected to all other ROIs) (Figure 1A). Then, we applied a hierarchical parametric empirical Bayes framework (Friston et al., 2016) to identify the subsets of excitatory and inhibitory connections associated with (i) homeostatic state (hunger versus satiety), (ii) obesity (BMI), (iii) homeostatic state-by-obesity interactions. We hypothesised that BMI would associate with increased influence of striatal and insula regions on the hypothalamus, and decreased influence of cortical regions on lower-level ones.

**Figure 1.**
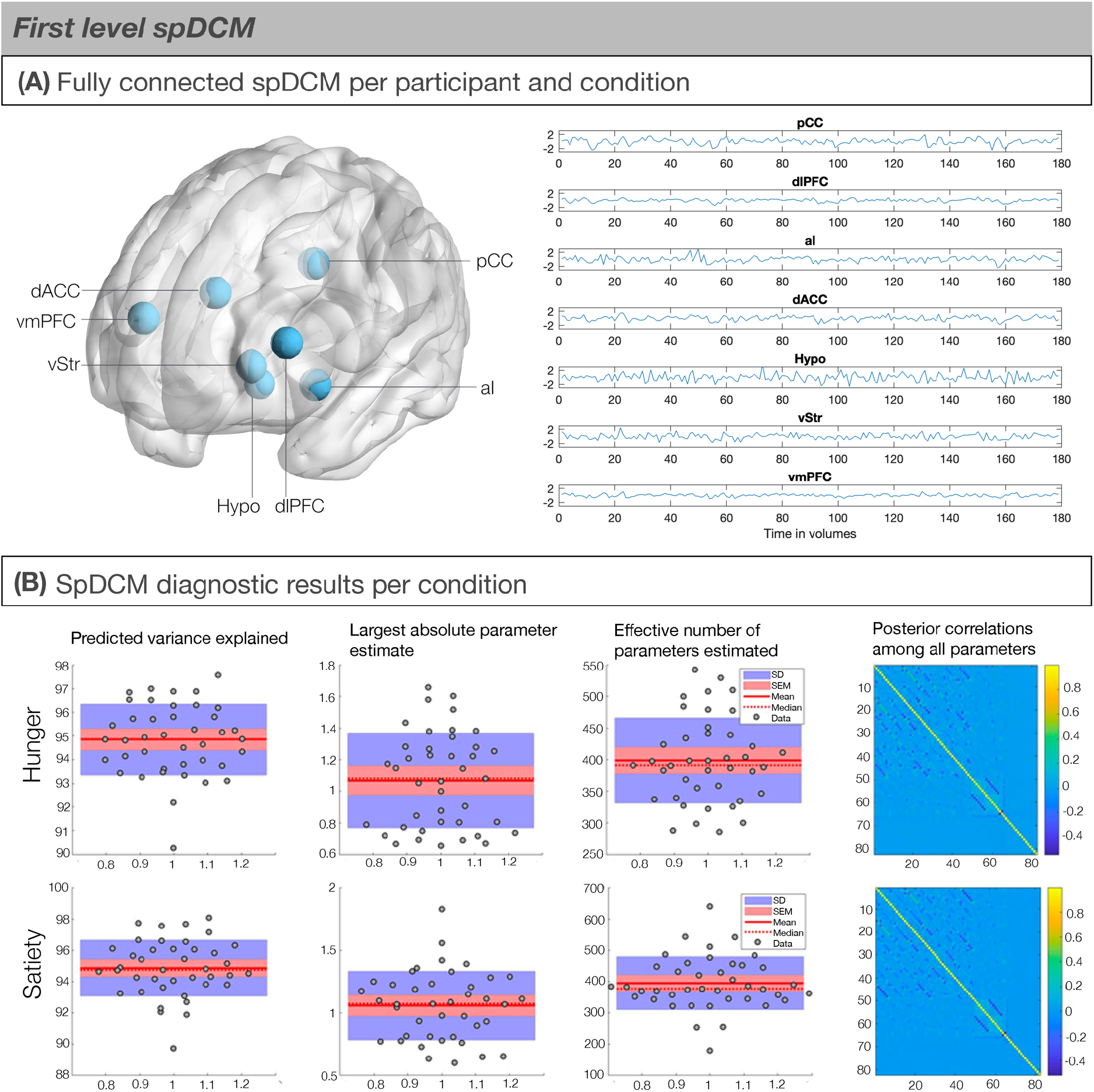
**(A)** Regions of interest (ROIs). ROI selection was based on task-based fMRI results within the same sample (Harding et al., 2018, Table 2). (Left) Schematic showing the seven ROIs used to estimate spDCMs with the fully-connected architecture (i.e., 7^2^ = 49 parameter model; Friston et al., 2014). (Right) The time series of the ROIs for an exemplar subject. Posterior cingulate cortex, pCC; Dorso-lateral prefrontal cortex, dlPFC; Anterior insula, aI; dorso-anterior cingulate cortex, dACC; Hypothalamus, Hypo; Ventral striatum, vStr; Ventromedial prefrontal cortex; vmPFC. **(B)** First level DCM model convergence statistics indicating good model convergence. (First column) Predicted variance explained for each individual were high mostly above 85%. (Second column) The largest absolute parameter estimate did not fall below the typical connection strength of 1/8Hz (Third column). The effective number of parameters are reported in terms of divergence between the posterior and prior densities over parameters. (Fourth or last column) Posterior correlations among all parameters were low, indicating identifiable parameters.

**Table 1.**
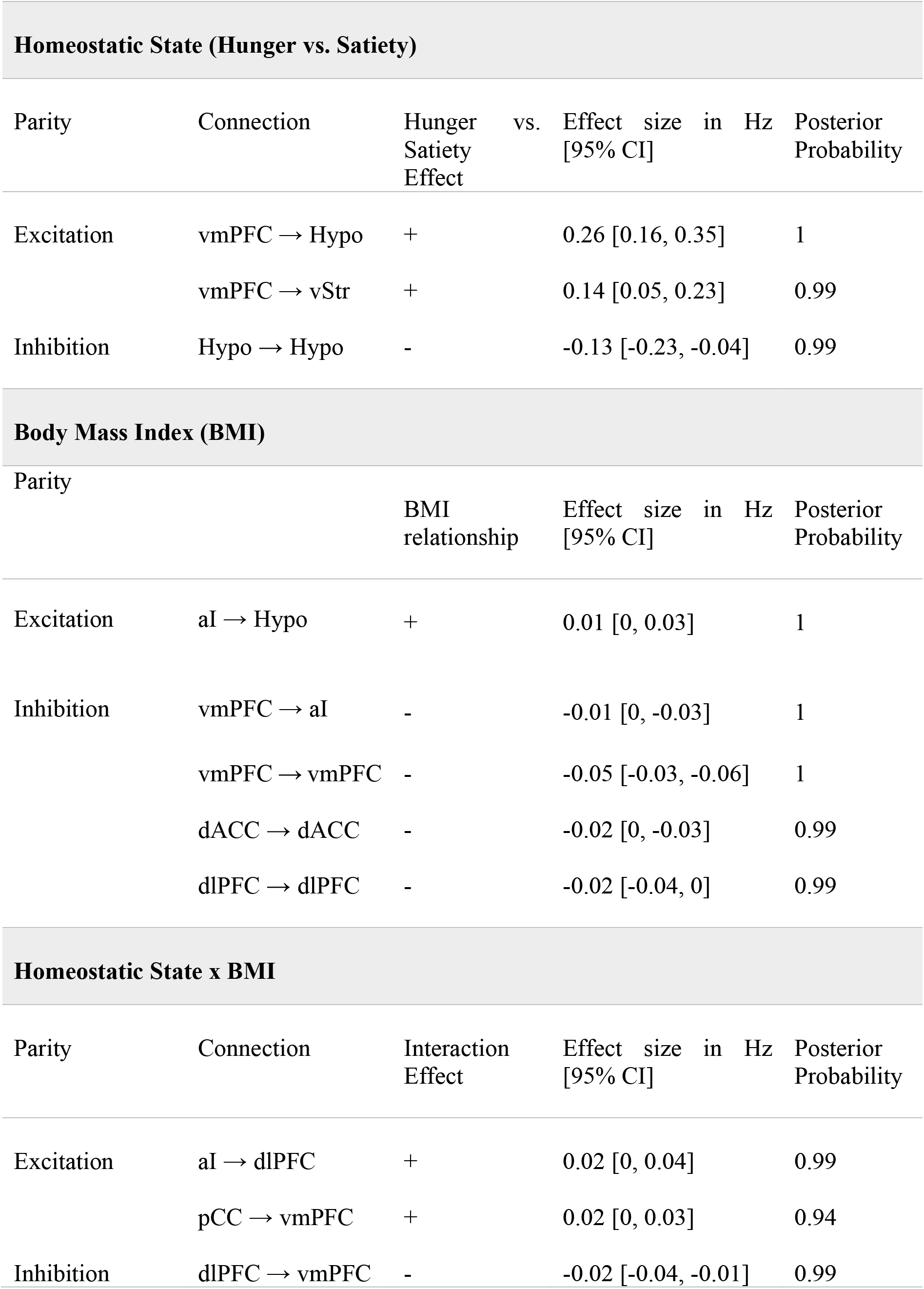
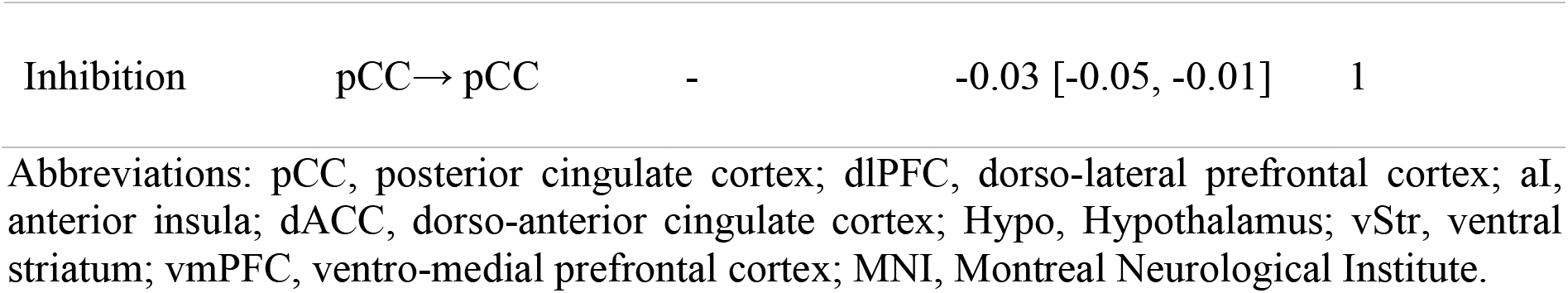
Summary of spDCM findings.

## Results

At the subject-level, we used the time-series from the ROIs to define and estimate a fully connected DCM for each participant and condition using Variational Laplace (Friston, Mattout, Trujillo-Barreto, Ashburner, & Penny, 2007) (Figure 1A). The average variance explained across subject-level DCM inversion was very high (Hunger: M = 94.85, SD = 1.50, range = 90.27-97.58; Satiety: M = 94.85, SD = 1.78, range = 89.73-98.06), indicating very good model fitting (Figure 1B).

### Homeostatic State (Hunger versus Satiety)

To explore how homeostatic state is associated with connectivity changes, we examined causal network dynamics during hunger and satiety while controlling for BMI. Starting from a fully-connected model (Figure 1), Bayesian optimisation procedures revealed a sparse model structure with a posterior probability of >.99 at the group level (Figure 2A, Table 1). Compared to satiety, hunger was associated with an increased excitatory influence of the ventromedial prefrontal cortex over the ventral striatum (0.14 Hz, 95% CI [0.05, 0.23]) and hypothalamus (0.26 Hz, 95% CI [0.16, 0.35]). We further found less self-inhibition (i.e., disinhibition) of the hypothalamus, which is thought to reflect an enhanced sensitivity to inputs from connecting nodes (Friston et al., 2003), when individuals were hungry as opposed to sated (−0.13 Hz, 95% CI [−0.23, −0.04]). Leave-one-out cross-validation revealed that these effects from individual connections are large enough to predict left-out individuals’ hunger state above chance level (r(df = 80) = 0.32, p < 0.002). Cross-validation of this sort provides out of sample estimates of predictability (i.e., the predictive validity of the connectivity strength from a new participant’s hunger state).

**Figure 2.**
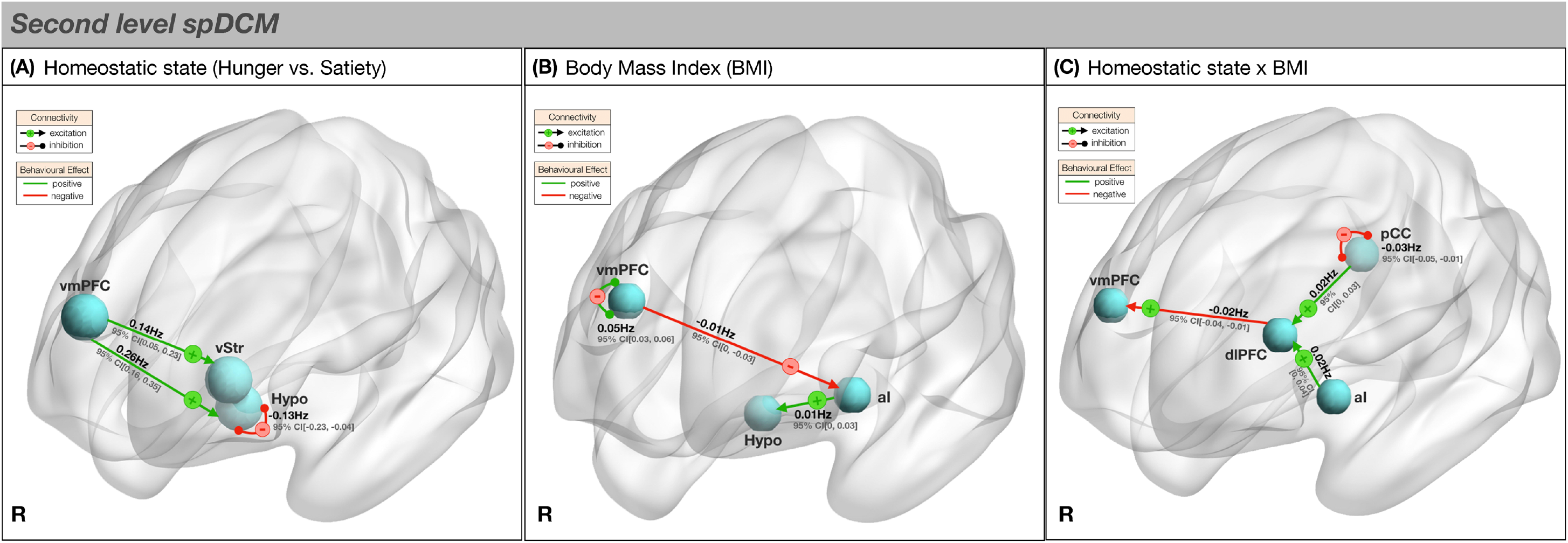
Second level spDCM results. **(A)** Effective connectivity of hunger. Hunger was associated with increased excitatory connectivity from the ventromedial prefrontal cortex to the ventral striatum and Hypothalamus and decreased hypothalamic self-inhibition. **(B)** Effective connectivity of BMI. BMI was associated with decreased inhibition from the ventromedial prefrontal cortex to the anterior insula, increased excitation from the anterior insula to the hypothalamus and increased self-inhibition of the ventromedial prefrontal cortex. **(C)** Effective connectivity of BMI x Hunger State. During hunger, compared to satiety, higher BMI was associated with decreased dorso-lateral prefrontal cortex to ventromedial prefrontal cortex excitation, increased excitation from the posterior cingulate cortex and anterior insula to the dorso-lateral prefrontal cortex and decreased posterior cingulate cortex self-inhibition. + or – signs code the parity of connectivity: –, inhibitory; +, excitatory. Table 1 provides further details of results. R, right, for all other abbreviations see text.

### Body mass index (BMI)

Next, we explored how BMI is associated with is associated with connectivity changes, whilst controlling for homeostatic state. Elevated BMI was associated with an increased excitatory influence of the anterior insula on the hypothalamus (0.01 Hz 95% CI [0, 0.03]) and a reduced inhibitory influence of the ventromedial prefrontal cortex on the anterior insula (−0.01 Hz, 95% CI [0, −0.03]) (Figure 2B; Table 1). In addition, individuals with greater BMI had increased self-inhibition of the ventromedial prefrontal cortex (0.05 Hz, 95% CI [0.03, 0.06]) and dorso-anterior cingulate cortex (0.02 Hz, 95% CI [0, 0.03]), and decreased self-inhibition of the dorso-lateral prefrontal cortex (−0.02 Hz, 95% CI [−0.04, 0]) and posterior cingulate cortex (−0.02 Hz, 95% CI [−0.03, 0]). Leave-one-out cross validation revealed that these effects sizes from individual connections are large enough to predict group effects with an out of sample estimate (r(df = 80) = 0.22, p < 0.05).

### Interaction of BMI and Homeostatic State

In the final analysis, we investigated how hunger-related connectivity changes may be associated with differences in BMI. During hunger relative to satiety, higher BMI was associated with a lesser excitatory influence of the dorso-lateral prefrontal cortex over the ventromedial prefrontal cortex (−0.02 Hz, 95% CI [−0.04, −0.01]) and a greater excitatory influence of the anterior insula over the dorso-lateral prefrontal cortex (0.02 Hz, 95% CI [0, 0.04]) (Figure 2C; Table 1). In addition, we found decreased self-inhibition of the posterior cingulate cortex (−0.03 Hz, 95% CI [−0.05, −0.01]). An increased excitatory influence of the posterior cingulate cortex on the dorso-lateral prefrontal cortex was also evident below the set posterior probability threshold of > 0.99 (0.02 Hz, 95% CI [0, 0.03], posterior probability = .94). The out of sample correlation between the model’s prediction and observed data was significant as revealed by leave-one-out cross validation (r(80)= 0.19, p < 0.05).

## Discussion

This study reveals novel obesity-related changes in directional interactions between corticostriatal and homeostatic regions (summarised in Figure 3). We specifically examined brain regions featured in neurobiological theories of obesity, including the hedonic eating, eating without hunger and self-regulation views. We found that higher BMI was associated with a greater excitatory influence of the anterior insula on the hypothalamus, regardless of homeostatic state (i.e. during both hunger and satiety). This finding is consistent with reduced sensitivity to changes in energy homeostasis and an eating without hunger view (Carnell et al., 2013; Carter et al., 2016; Volkow et al., 2017). Furthermore, participants with higher BMI showed weaker excitatory influence of the dorso-lateral prefrontal cortex on the ventromedial prefrontal cortex during the hunger state. The interaction between these two regions have been previously associated with dietary self-regulation (Hare et al., 2009). In addition, we showed that, regardless of adiposity, during hunger as compared to satiety the ventromedial prefrontal cortex increased its excitatory influence over the ventral striatum and the hypothalamus. This may represent a general adaptive mechanism of top-down signalling during energy deprivation (Morton et al., 2014).

**Figure 3.**
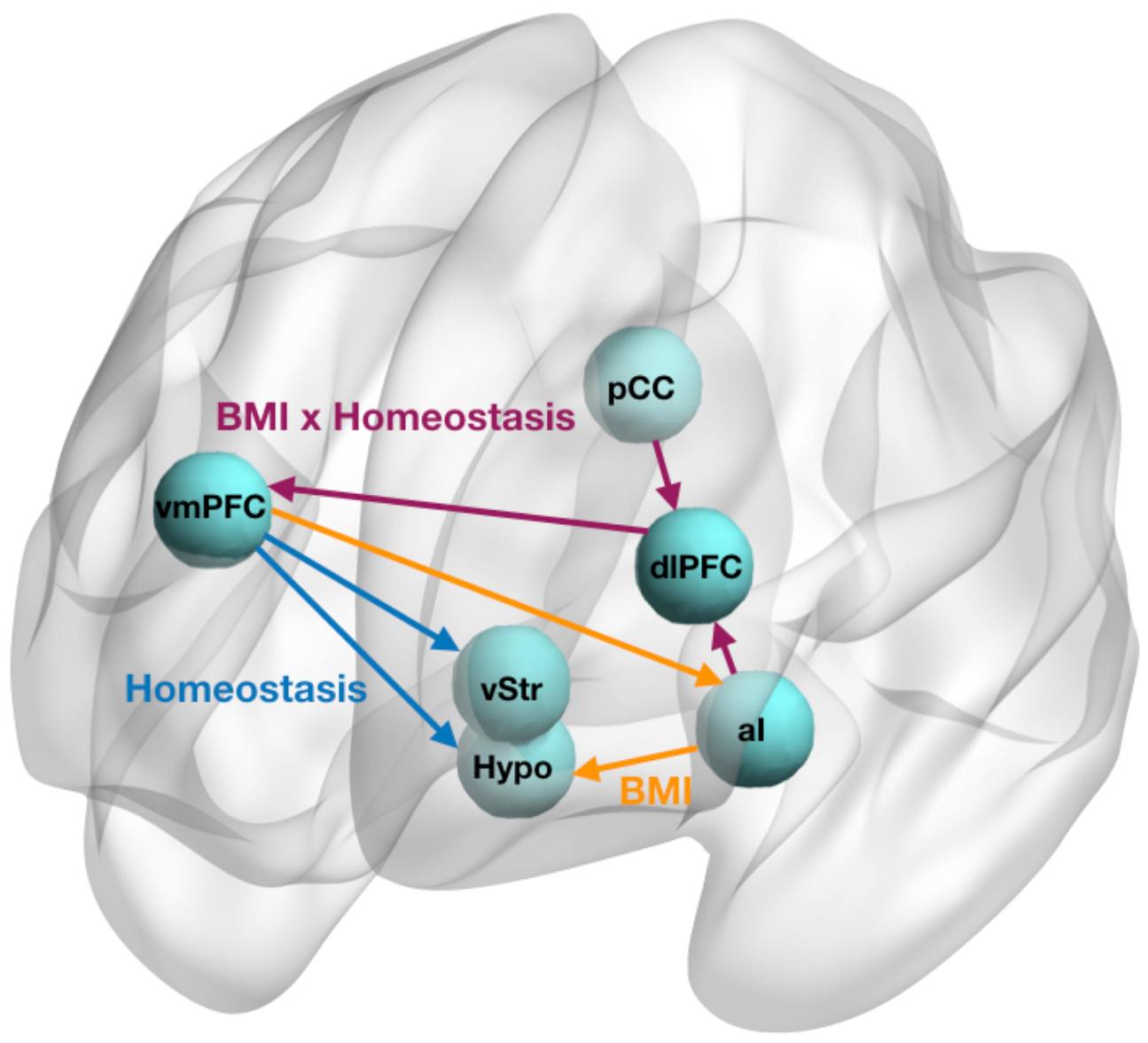
Schematic diagram summarising the resting state neuronal network configurations associated with Homeostasis, BMI and Homeostasis x BMI interaction effects.

Together with the hypothalamus, the anterior insula has been proposed to form a homeostatic / interoceptive network that prompts eating during energy deprivation and ends feeding upon satiation in humans (Wright et al., 2016) and plays a central role in food-seeking behaviour in rodents (Kusumoto-Yoshida et al., 2015). Our findings suggest that this mechanism is altered in obesity, as increased excitatory input from the insula to the hypothalamus persists during satiety. This occurs together with reduced inhibition from the ventromedial prefrontal cortex to the anterior insula - a pathway that codes changes in incentive salience in response to changes in homeostatic state (Damasio et al., 1991; Verdejo-García and Bechara, 2009). In fitting with this interpretation, obesity would be associated with reduced neural sensitivity to changes in homeostatic / interoceptive state and related persistent attribution of salience to both hunger and satiety states. This argument is consistent with the eating without hunger framework (Carnell et al., 2013).

Furthermore, we found that obesity was associated with changes in cortico-cortical interactions during the hunger state. Reductions in dorso-lateral prefrontal cortex influence over the ventromedial prefrontal cortex, as observed herein, play a central role in goal-directed food choice tasks and related dietary self-regulation (Hare, Malmaud, & Rangel, 2011; Hare et al., 2009). Although we cannot assume equivalence between the function of brain regions in task-related versus resting-state designs (Poldrack, 2006; Jung et al., 2018), since these regions were activated by a food choice task in the same participants (Harding et al., 2018), and in absence of more plausible alternative explanations, we speculate that these findings may relate to alterations in goal-oriented food choice. If our interpretation is correct, these results would align with an impaired self-regulation model of obesity (Carter et al., 2016; Volkow et al., 2017) but introduce the additional caveat that this mechanism may be state-specific as it was not observed in the satiety state. The obesity-related greater excitation from anterior insula and posterior cingulate regions over the dorso-lateral prefrontal cortex is consistent with this hunger-related effect (Al-Zubaidi et al., 2019).

Communication between “higher-order” cortical regions and “lower-order” subcortical areas (e.g., hypothalamus) are critical to governing feeding behavior (Andermann and Lowell, 2017). Our results in the hunger (versus satiety) state, irrespective of BMI, support a top-down interpretation of these relationships, in line with the notion that hunger triggers an incentive mechanism during energy depletion that motivates food seeking in order to avoid starvation (Krashes et al., 2011; Betley et al., 2015). Studies in rodent and primate models similarly support the role of ventromedial prefrontal cortex-hypothalamus interactions in feeding behavior (Ongur and Price, 2000). Notably, we previously reported that homeostatic state influences local activity with the hypothalamus itself (Harding et al., 2018), in line with its established sensitivity to the neuropeptides that signal and regulate energy needs (e.g. ghrelin and leptin; (Farooqi et al., 2007; Malik et al., 2008).

In conclusion, our study is the first to provide changes in directed connectivity between prefrontal, insular, striatal and hypothalamic regions as a function of BMI and homeostatic state. Although the hedonic eating view has been a dominant account of obesity, our findings point to a model in which reduced sensitivity to homeostatic / interoceptive changes and disrupted prefrontal communication during caloric deprivation. Our results highlight the potential for intrinsic predispositions (“neuromarkers”), to improve individualised intervention strategies that can either alter obesogenic traits or ameliorate detrimental effects such as weight gain. They also pave the way for developing non-invasive brain stimulation protocols aimed to rewire cortico-subcortical network abnormalities in obesity. Specifically, our findings provide proof-of-principle evidence to examine the impact of dorso-lateral prefrontal cortex stimulation and ventromedial prefrontal cortex inhibition on cortical-insular-hypothalamic network dynamics and obesity-related therapeutic outcomes. While our study is only the first step to deriving an understanding of this interplay of brain regions at rest and how this might translate into the expression of the behavioural effects, complementary future task-based studies will be required to examine how the revealed resting-state network dynamics translate during task performance.

## Methods

### Participants

Forty-one participants (21 females, mean age = 24.37, SD = 5.53) were recruited from the general community. Participants’ BMI ranged from 18 to 38kg/m^2^: 20 had healthy weight (18-24.9kg/m^2^), 21 were overweight (BMI = 25-30 kg/m^2^) or obese (BMI > 30 kg/m^2^). An initial screening interview assured that these participants (1) had no history of hypertension or diabetes, (2) had no neurological and psychiatric illness, or (3) were on psychoactive medication affecting cognitive functioning or cerebral blood flow. All participants were naive to the purpose of the study, gave written consent before participating, and were reimbursed with $100 gift card vouchers. The Monash University Human Research Ethics Committee approved the study (2019-5979-30222) following the Declaration of Helsinki.

### Experimental procedure and MRI data acquisition

Participants completed two resting-state fMRI scans, one after an overnight fast (hunger condition) and one after a standard breakfast (satiety condition). In both conditions, participants were instructed to have a standard meal (700-1000kj) between 7.30pm and 8.30pm on the night prior to their scan refrain from eating or drinking (except for water) until their morning scan. For the satiety condition, participants received a standard breakfast (293 kcal) one hour prior to their scan. Subjective self-reports of hunger (1 = not hungry at all; 7 = very hungry) revealed a significant difference in the perception of hunger between hunger (M = 4.63; SD = 1.46) and satiety (M = 3.12; SD = 1.58) condition (t(40) = 4.72, p < 0.001). There was no difference in subjective feelings of fullness across the weight groups when they received a meal, therefore we did not include these parametric regressors into our main spDCM analyses. All scans were scheduled in the morning between 9am and 10am and on average there were 5.82 days (SD = 3.73 days) between the two scanning sessions. Scheduling the participant’s first scan as either part of the hunger or satiety condition was counterbalanced across participants.

Resting-state fMRI data were acquired using a 3-Tesla Siemens Skyra MRI scanner equipped with a 32-channel head coil at the Monash Biomedical Imaging Research Centre (Melbourne, Victoria, Australia). During each 8-minute scan, 189 gradient-echo planar images comprising of 44 interleaved, continuous axial slices were collected (repetition time = 2500 ms; echo time = 30 ms; flip angle = 90°; 3 mm isotropic voxels; field of view = 192 mm). A whole-brain T1-weighted magnetisation-prepared rapid gradient-echo structural image was also acquired for each session and each participant (192 sagittal slices; 1 mm isotropic voxels; repletion time = 2300 ms; field of view = 256 mm). Participants were instructed to rest while closing their eyes.

Following the resting-state fMRI scan of each session, participants completed a task-based fMRI session, during which they were required to make realistic choices between unhealthy and healthy drinks. ROI selection was based on the results revealed during the task-based fMRI session (Table 2). The full study protocol and results are reported elsewhere (Harding et al., 2018). The task-based session was always performed after the resting-state MRI session, to avoid confounding the resting-state BOLD signals (Tambini et al., 2009).

**Table 2.** Locations of regions of interests (spDCM nodes).

| Region | MNI Coordinates |  |  | Source |
| --- | --- | --- | --- | --- |
|  | x | y | z |  |
| pCC | 2 | -42 | 30 | Harding et al. (2018) |
| dIPFC | -45 | 42 | 16 | Harding et al. (2018) |
| aI | -38 | 12 | -6 | Harding et al. (2018) |
| dACC | -4 | 32 | 30 | Harding et al. (2018) |
| Hypo | 0 | -3 | -12 | Neurosynth |
| vStr | -4 | 8 | -3 | Batra et al. (2013) |
| vmPFC | 12 | 58 | 22 | Harding et al. (2018) |

Abbreviations: pCC, posterior cingulate cortex; dIPFC, dorso-lateral prefrontal cortex; aI, anterior insula; dACC, dorso-anterior cingulate cortex; Hypo, Hypothalamus; vStr, ventral striatum; vmPFC, ventro-medial prefrontal cortex; MNI, Montreal Neurological Institute.

### Resting-state fMRI data analyses

#### Preprocessing

Functional images were preprocessed using SPM12 (revision 12.2, www.fil.ion.ucl.ac.uk). Following preprocessing guidelines for spDCM (Park et al., 2018), the preprocessing steps consisted of slice time correction, realignment, spatial segmentation and normalisation to the standard EPI template of the Montreal Neurological Institute (MNI), and spatial smoothing using a Gaussian kernel of 8-mm FWHM. No (band-pass) filtering was used except a low-pass filter (of 1/128) that filters the ultra-low frequency scanner drifts (Razi et al., 2015). None of the participants exceeded excessive head motion of larger than 3mm.

#### ROI selection and time series extraction

Seven ROIs were identified as key nodes for effective connectivity analyses. The identified neural circuit comprised of the posterior cingulate cortex, dorso-lateral prefrontal cortex, anterior insula, dorso-anterior cingulate cortex, Hypothalamus, ventral striatum, and ventromedial prefrontal cortex (Figure 1A). The MNI coordinates for these regions were based on activity associated with food choices during the task-based fMRI session (Harding et al., 2018; Table 2).

To extract BOLD fMRI time series corresponding to the aforementioned ROIs, the pre-processed data was used to establish the residuals of a General Linear Model (GLM). Six head motion parameters and WM/CSF signals were added to the GLM as nuisance regressors. Finally, we selected the MNI coordinates as the centre of a 6-mm sphere to compute the subject-specific principal eigenvariate and correct for confounds.

### Spectral Dynamic Causal Modeling

The spDCM analyses were performed using the functions of DCM12 (revision 7196) implemented in SPM12. In order to address our main hypotheses, we focused on spDCM analyses that assessed (1) changes in effective connectivity of hunger versus satiety condition independent of BMI (main effect of hunger), (2) changes in effective connectivity modulated by BMI (main effect of BMI), and (3) changes in hunger-related effective connectivity modulated by BMI (BMI-by-hunger interaction).

#### First Level spDCM Analysis

On the first-level, a fully-connected model was created for each participant and each session (i.e. 7^2^ = 49 connectivity parameters, including seven inhibitory self-connections). Next, we inverted (i.e. estimated) the DCMs using spectral DCM, which fits the complex cross-spectral density using a parameterised power-law model of endogenous neural fluctuations (Razi et al., 2015, 2015). This analysis provides measures of causal interactions between regions, as well as the amplitude and exponent of endogenous neural fluctuations within each region (Razi et al., 2015). Model inversion was based on standard variational Laplace procedures (Friston et al., 2007). This Bayesian inference method uses Free Energy as a proxy for (log) model evidence, while optimising the posterior density under Laplace approximation of model parameters.

#### Second Level spDCM Analysis

To characterize how group differences in neural circuitry were modulated by BMI and Hunger condition, hierarchical models over the parameters were specified within a hierarchical Parametric Empirical (PEB) framework for DCM (Friston et al., 2016)

##### Model 1: Effective connectivity of Homeostatic State

To investigate the main effect of homeostasis (i.e., hunger condition as opposed to satiety condition) on the intrinsic effective connectivity of the neural network of interest, the following hierarchical model was estimated within a PEB framework:

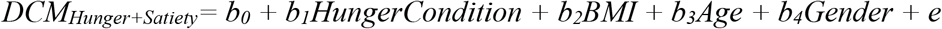

Hunger condition was modelled as the main regressor of interest as a vector consisting of 1 (fasted) and −1 (stated). Mean-centered BMI, Age and Gender where modelled as regressors of no interests.

##### Model 2: Effective connectivity of BMI

To investigate the main effect of increased BMI on the causal dynamics within the ROIs network, the following hierarchical model was estimated within a PEB framework:

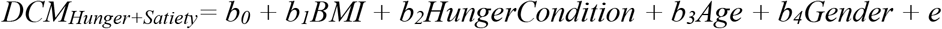

BMI was modelled as the main regressor of interest, whereas Hunger Condition, Age and Gender where modelled as regressors of no interests.

##### Model 3: Effective connectivity of Homeostatic State x BMI interaction effect

Model 3 was designed to assess how the interaction between BMI and Hunger Condition affects the dynamics of the network.

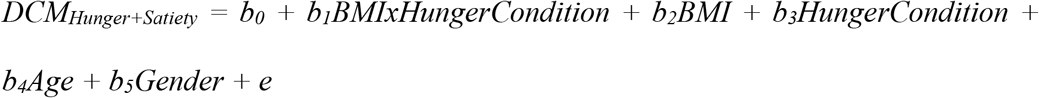

The interaction term between BMI and Hunger Condition was created by first centering the continuous variable BMI before creating the element-by-element product of the newly centered BMI variable with the categorical variable Hunger condition (Aiken and West, 1991). The interaction effect was modelled as main regressor of interest in Model 2, whereas the main effects of BMI (centered), Hunger Condition, Age, Gender were regressors of no interest.

Note that for each of the presented models, all behavioural regressors were mean-centered so that the intercept of each model was interpretable as the mean connectivity. Bayesian model reduction was used to test all reduced models within each parent PEB model (assuming that a different combination of connections could exist (Friston et al., 2016) and ‘pruning’ redundant model parameters; parameters of the best 256 pruned models (in the last Occam’s window) were averaged and weighted by their evidence (i.e. Bayesian Model Averaging) to generate final estimates of connection parameters. To identify important effects (i.e., changes in directed connectivity), we compared models (using log Bayesian model evidence to ensure the optimal balance between model complexity and accuracy) with and without each effect and calculated the posterior probability for each model, as a softmax function of the log Bayes factor. We treat effects (i.e., connection strengths and their changes) with posterior probability >0.99 as significant for reporting purposes.

Finally, in order to determine the predictive validity (e.g. whether BMI can be predicted from the final, reduced spDCM’s individual connections), leave-one-out cross validation was performed within the PEB framework (Zeidman et al., 2019). This procedure fits the PEB model to all but one participant and predicts the covariate of interest (e.g. BMI) for the left-out participant. This is repeated with each participant to assess the averaged prediction accuracy for each model.

## Acknowledgements

The authors thank Richard McIntyre for help with MRI data acquisition. This study was supported by a NHMRC grant (1140197) granted to Antonio Verdejo-Garcia.

